# Notch3 Promotes Prostate Cancer-Induced Bone Lesion Development via MMP-3

**DOI:** 10.1101/349043

**Authors:** Sourik S. Ganguly, Galen Hostetter, Lin Tang, Sander B. Frank, Kathylynn Saboda, Rohit Mehra, Lisha Wang, Xiaohong Li, Evan T. Keller, Cindy K. Miranti

**Affiliations:** Program for Skeletal Disease and Tumor Microenvironment, Center for Cancer and Cell Biology; Tissue Repository and Pathology Core, Van Andel Research Institute, Grand Rapids, MI, USA; Department of Cellular and Molecular Medicine, University of Arizona Cancer Center; University of Arizona Cancer Center, University of Arizona, Tucson, AZ, USA; Department of Pathology, University of Michigan, Ann Arbor, MI, USA

**Author notes:** Corresponding Author: Cindy K Miranti, University of Arizona Cancer Center, 1515 N. Campbell Ave, Tucson, AZ 85718.

**Keywords:** prostate cancer bone metastasis, notch, MMP, osteoblasts, osteoclasts

## Abstract

Prostate cancer metastases primarily localize in the bone where they induce a unique osteoblastic response. Elevated Notch activity is associated with high-grade disease and metastasis. To address how Notch affects prostate cancer bone lesions, we manipulated Notch expression in mouse tibia xenografts and monitored tumor growth, lesion phenotype, and the bone microenvironment. Prostate cancer cell lines that induce mixed osteoblastic lesions in bone expressed 5-6 times more Notch3, than tumor cells that produce osteolytic lesions. Expression of active Notch3 (NICD3) in osteolytic tumors reduced osteolytic lesion area and enhanced osteoblastogenesis, while loss of Notch3 in osteoblastic tumors enhanced osteolytic lesion area and decreased osteoblastogensis. This was accompanied by a respective decrease and increase in the number of active osteoclasts and osteoblasts at the tumor-bone interface, without any effect on tumor proliferation. Conditioned medium from NICD3-expressing cells enhanced osteoblast differentiation and proliferation in vitro, while simultaneously inhibiting osteoclastogenesis. MMP-3 was specifically elevated and secreted by NICD3-expressing tumors, and inhibition of MMP-3 rescued the NICD3-induced osteoblastic phenotypes. Clinical osteoblastic bone metastasis samples had higher levels of Notch3 and MMP-3 compared to patient matched visceral metastases or osteolytic metastasis samples. We identified a Notch3-MMP-3 axis in human prostate cancer bone metastases that contributes to osteoblastic lesion formation by blocking osteoclast differentiation, while also contributing to osteoblastogenesis. These studies define a new role for Notch3 in manipulating the tumor microenvironment in bone metastases.

## INTRODUCTION

Death from prostate cancer is primarily due to metastasis (1, 2). Prostate cancer invariably metastasizes to the bone and forms osteoblastic (bone-forming) lesions. Such lesions cause severe bone pain, fractures, bone deformity, hypercalcemia, and immunological complications (1, 3). Even though prostate cancer metastasis is typically osteoblastic, markers of bone turnover indicate that underlying osteolytic events are also involved (4, 5). Death from metastatic bone disease is due in part to the lack of effective therapy, the development of which requires knowing the mechanisms that control osteoblastic versus osteolytic phenotypes.

Bone is constantly remodeling during an individual’s lifetime. Remodeling depends on two cells types, osteoblasts and osteoclasts, which work in harmony to maintain normal bone. Osteoblasts, derived from mesenchymal stem cells, make new bone. Osteoclasts, derived from monocytes, degrade bone. During osteoblastic metastasis, the bone remodeling balance favors bone formation, i.e. osteoblastogenesis (1, 6).

Notch, best known for its role in development and differentiation, is also involved in cancer progression and metastasis. There are four single-pass transmembrane Notch receptors in mammals, activated by five different transmembrane ligands (JAG1/2 and DLL1/3/4) from adjacent cells upon cell-cell contact. Upon ligand binding, Notch is first cleaved by ADAM10 and then by γ-secretase to release the Notch intracellular cytoplasmic domain (NICD). NICD translocates to the nucleus and forms a transcription activation complex by associating with the proteins CBF-1, Suppressor of Hairless, Lag2, and Mastermind-like (MAML). This complex activates the transcription of Notch target genes like Hairy enhancer of split (Hes) and Hes-related YRPF motif (Hes) transcription factors (7–9). Majority of the studies investigating normal prostate gland development and prostate cancer have focused on Notch1. In the normal prostate, Notch1 is required for basal cell maintenance, luminal progenitor proliferation, and branching morphogenesis (10, 11). In prostate cancer, knockdown of Notch1 inhibits invasion, decreases growth, suppresses colony formation *in vitro*, and increases chemosensitivity (12, 13). Elevated Notch1 staining is observed in aggressive prostate tumors relative to prostatic intraepithelial neoplasia and normal tissue (12). These data suggest Notch1 may be a driver of prostate cancer progression.

Recent studies demonstrate that Notch3 is involved in normal prostate development where it is required for luminal cell differentiation (11, 14, 15). Furthermore, elevated Notch3 expression positively correlates with prostate cancer progression, recurrence, and drug resistance (16–20). In metastatic breast cancer, Jag-1 in tumor cells activates Notch signaling in osteoblasts to stimulate osteoclastogenesis in osteolytic bone lesions, while Notch3 is induced in tumor cells by osteoblasts (21, 22). However, the role of Notch3 in cancer cells and its influence on osteolytic versus osteoblastic disease remains unknown.

Matrix metalloproteinases (MMPs) are a family of proteases secreted by stromal and tumor cells which increase in many cancers and negatively correlate with patient survival (23). MMP-1, 3, 7 and 9 reportedly promote prostate cancer metastasis to the bone (5). Both MMP mRNA and protein are elevated in serum and tissue samples from prostate cancer patients and is correlated with advanced and metastatic disease (23). Polymorphisms in MMP-3 predict for increased prostate cancer risk and MMP-3 levels are elevated specifically in patients with bone metastases (24, 25). MMP-1, 9, and 13 are required for bone resorption, and MMP-9 promotes osteoclast polarization and ruffled border formation (5). Although Notch1 reportedly promotes the expression of MMP-9 (12), it is unknown whether Notch3 contributes to the expression of MMPs.

During our studies, we observed elevated Notch3 expression in human prostate cancer bone metastases and hypothesized that Notch3 promotes bone metastasis by remodeling the bone microenvironment. Herein, we report that Notch3 contributes to osteoblastic bone metastasis by suppressing osteoclasts and stimulating osteoblastogenesis through induction of MMP-3.

## RESULTS

### Notch3 inhibits osteolytic lesion development

Prostate cancer cells, which promote osteolytic (PC3) or mixed (blastic/lytic) lesions (C4-2B and 22Rv1), were assayed for Notch1 and Notch3 expression by qRT-PCR and immunoblotting. Both Notch1 and Notch3 mRNAs were significantly higher in the osteoblastic lines, C4-2B and 22Rv1, relative to the lytic PC3 cells (Fig. 1A-B). Notch1 and Notch3 protein were similarly higher in osteoblastic lines (Fig. 1C-D) suggesting Notch might play a role in osteoblastic lesion development.

**Figure 1.**
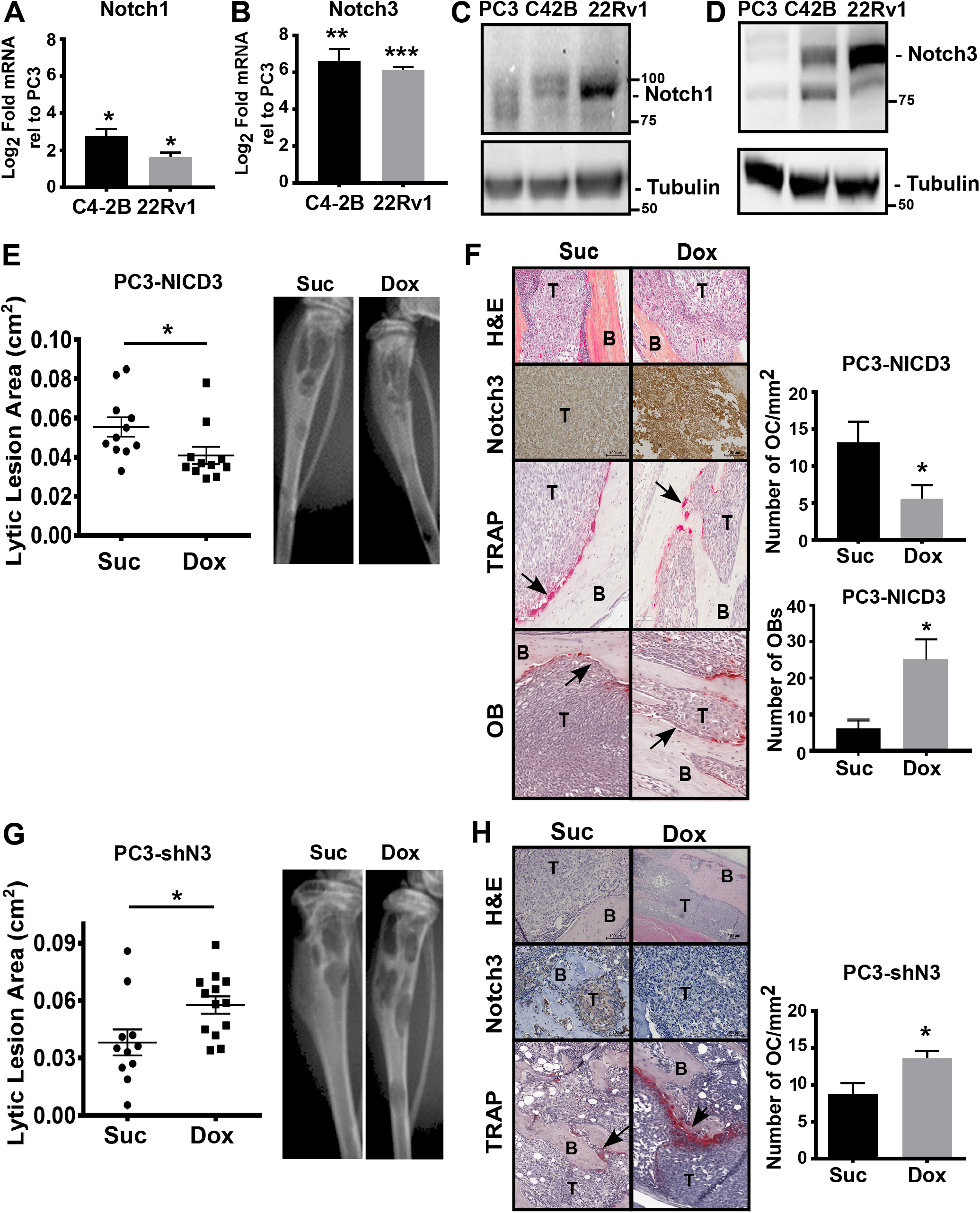
NICD3 inhibits osteolytic lesions. Levels of **(A)** Notch1 and **(B)** Notch3 mRNA in tumor cell lines. Log_2_-fold change relative to PC3 cells. Levels of **(C)** Notch1 and **(D)** Notch3 measured by immunoblotting. Tubulin is loading control. **(E,G)** PC3 cells harboring Tet-inducible **(E)** NICD3 or **(G)** Notch3 shRNA (shN3) injected into tibiae of mice treated with sucrose (Suc) or doxycycline (Dox). X-rayed lytic lesion area quantified. **(F,H)** Tibiae from (E) and (G) were stained with H&E (top, bottom panel of F), anti-Notch3 (2nd panel), TRAP (3rd panel), and number of osteoclasts and osteoblasts (OB) quantified. T=tumor; B=bone; arrows indicate examples of an osteoclast or osteoblast. Error bars are S.E.M, n>11; *0.01≤p≤0.05; **0.001≤p<0.01; ***p<0.001.

Osteolytic PC3 cells were engineered to overexpress doxycycline-inducible Notch3 intracellular domain, NICD3 (Supplementary Fig. S1A). Mice with PC3-NICD3 cells injected into the tibiae were fed sucrose as controls or doxycycline (Dox) to induce NICD3 expression. Radiographic imaging after 3 weeks showed the expected osteolytic lesion development in control PC3-NICD3-injected tibiae (Fig. 1E). Induction of NICD3 by Dox significantly reduced (1.5-fold) the osteolytic lesion area. H&E staining validated tumor growth in the bone, and Notch3 IHC staining confirmed NICD3 induction in the tumors of Dox-treated mice (Fig. 1F).

PC3 cells engineered to induce NICD1 (Supplementary Fig. S1B), did not produce any significant changes in osteolytic lesion area when tibia-injected tumors were stimulated with Dox (Supplementary Fig. S2A). The observed changes in lesion area were not due to non-specific effects of Dox, as Dox treatment of mice injected with non-NICD parental expressing tumors did not alter osteolytic lesion formation (Supplementary Fig. S2B). Thus, specific induction of NICD3, but not NICD1, reduces osteolytic lesion formation in prostate tumors in the bone.

To determine whether the decrease in osteolytic lesion development by NICD3 was due to changes in tumor proliferation, we monitored Ki67 expression by IHC staining. There was no difference in proliferation between PC3 control and PC3 NICD3-expressing tumors (Supplementary Fig. S3A). There were also no changes in tumor cell proliferation detected when PC3-NICD3 cells were treated with Dox in culture (Supplementary Fig. S4A). To assess the effects of Notch3 on the bone microenvironment, we quantified the number of TRAP-positive osteoclasts. There was over a 2-fold decrease in the number of TRAP-positive osteoclasts at the tumor-bone interface in PC3-NICD3 expressing tumors compared to control tumors (Fig. 1F). Conversely, NICD3 increased 5-fold the number of osteoblasts (OB) at the bone-tumor interface (Fig. 1F). Thus, the effect of NICD3 is on the tumor microenvironment, rather than an intrinsic effect on tumor growth.

### Inhibiting Notch3 increases osteolytic lesions

To determine if loss of Notch3 enhances osteolysis, we injected PC3 cells harboring doxycycline-inducible shNotch3 (PC3-shN3) into mouse tibiae. PC3-shN3 Dox-treated tumors had significantly more lytic lesion area (1.5-fold) as compared to sucrose-treated control tumors (Fig. 1G). Loss of Notch3 expression was validated by immunoblotting after treating PC3-shNotch3 expressing cells with Dox (Supplementary Fig. S1D) and by IHC of tumors from mouse tibiae (Fig. 1H). There was no change in tumor proliferation (Supplementary Fig. S3B) and the Dox-treated PC3-shN3 tumors had significantly more TRAP positive cells (Fig. 1H). These results indicate that inhibiting Notch3, even in low Notch3 expressing cells, further promotes lytic lesion formation.

To assess whether blocking Notch3 expression in osteoblastic tumors enhances lytic lesions, we injected 22Rv1 or C4-2B cells harboring doxycycline-inducible shNotch3 into mouse tibiae. Loss of Notch3 in Dox-treated 22Rv1 and C4-2B tumors significantly increased osteolytic lesion area by 2.2-fold and 1.3-fold respectively (Fig. 2A,C). Loss of Notch3 expression was confirmed by immunoblotting after treating 22Rv1 and C4-2B shNotch3 cells with Dox (Supplementary Fig. S1E-F) and by IHC of tumors from mouse tibiae (Fig. 2B,D). Loss of Notch3 also resulted in a 2-fold increase in the number of osteoclasts at the tumor-bone interface (Fig. 2E). Conversely, loss of Notch3 decreased 5-fold the number of osteoblasts at the bone-tumor interface (Fig. 2F). Loss of Notch3 had no effect on tumor proliferation *in vivo* (Supplementary Fig. S3C-D) and there were also no changes in tumor cell proliferation when 22Rv1 and C4-2B cells were treated with Dox in vitro (Supplementary Fig. S4B-C). Thus, expression of active Notch3 in prostate tumors in the bone inhibits osteolytic lesion development independent of tumor proliferation.

**Figure 2.**
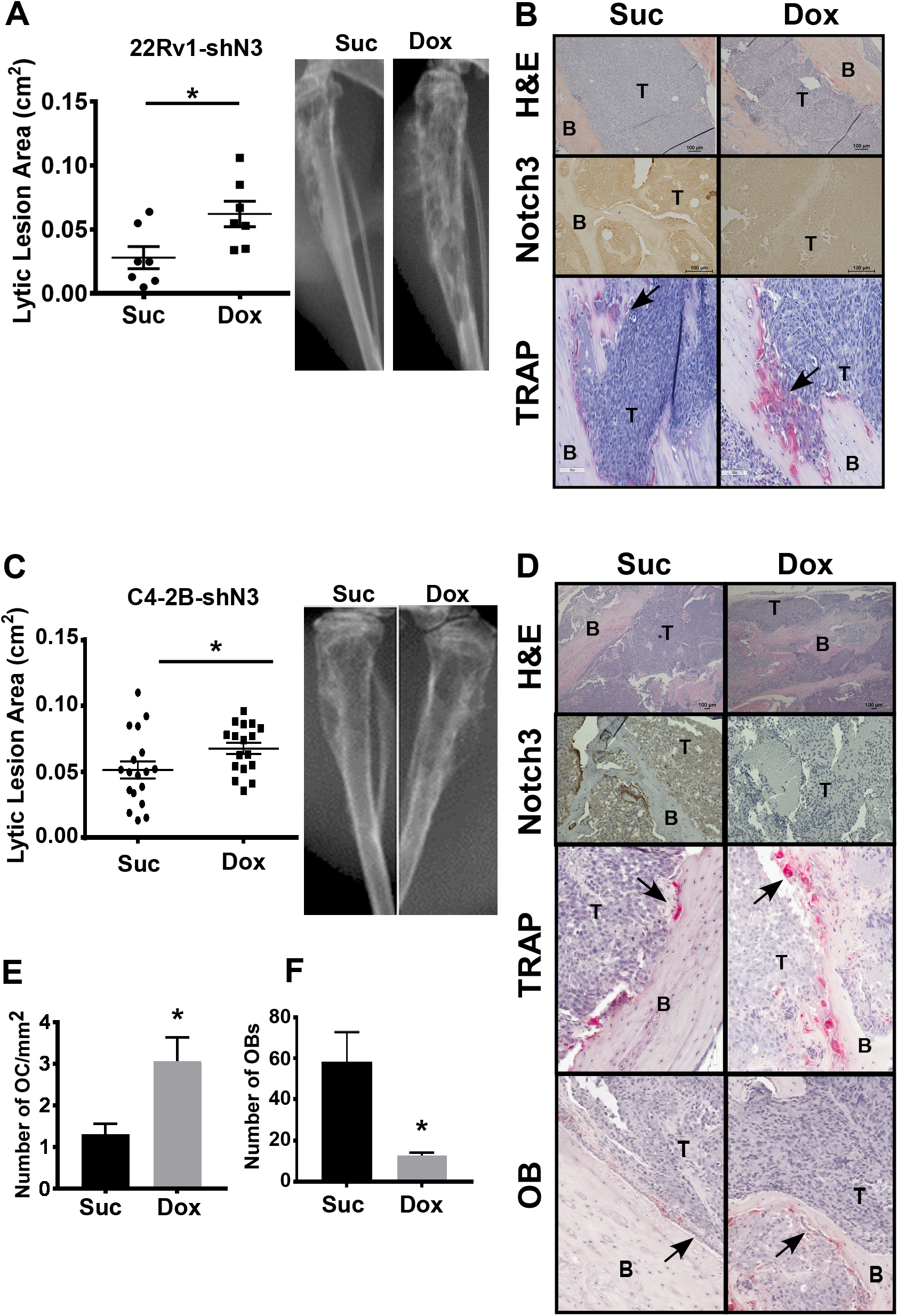
Inhibiting Notch3 promotes osteolytic lesions. **(A-B)** 22Rv1 or **(C-F)** C4-2B cells harboring doxycycline-induced Notch3 shRNA (shN3) injected into tibiae of mice treated with sucrose (Suc) or doxycycline (Dox). **(A, C)** X-rayed lytic lesion area quantified. **(B, D)** Tibiae were stained with H & E (top, bottom panel of D), anti-Notch3 (2nd panel), TRAP (3rd panel). **(E)** Number of osteoclasts quantified. **(F)** Number of osteoblasts (OB) quantified. T=tumor; B=bone; arrows indicate examples of an osteoclast or osteoblast. Error bars are S.E.M, n>7; *0.01≤p≤0.05.

### Notch3 induces the expression of MMP-3

To identify secreted factors that contribute specifically to osteoblastic bone lesion development, cell lysates from dissected subcutaneous tumors or crushed tibia from PC3, C4-2B, or 22Rv1 cells were applied to a cytokine array. MMP-3 was specifically elevated in C4-2B and 22Rv1 bones, compared to subcutaneous tumors (Supplementary Fig. S5). No change in MMP-3 levels was observed between PC3 tumors in the bone versus skin.

To investigate whether Notch3 controls MMP-3 expression, PC3-NICD3, PC3-shN3, or C4-2B-shN3 tibia-injected tumors treated with or without Dox were analyzed for MMP-3 expression by IHC and qRT-PCR. MMP-3 protein was elevated in the Dox-treated PC3-NICD3 tumors relative to sucrose-treated control tumors (Fig. 3A). Conversely, loss of Notch3 expression in PC3-shN3 or C4-2B-shN3 tumors by Dox reduced MMP-3 levels relative to sucrose-treated tumors (Fig. 3A). Similarly, human-specific MMP-3 mRNA was increased by Dox in PC3-NICD3 tumors (Fig. 3B), and decreased by Notch3 loss in PC3-shN3 and C4-2B-shN3 tumors (Fig. 3C-D). Furthermore, induction of NICD3 in PC3 cells in culture induced the secretion of pro and active MMP-3 into the conditioned medium (Fig. 3E), indicating NICD3 induces the expression of MMP-3. Levels of secreted pro and active MMP-3 ranged from 0.5ng/ml to 1ng/ml.

**Figure 3.**
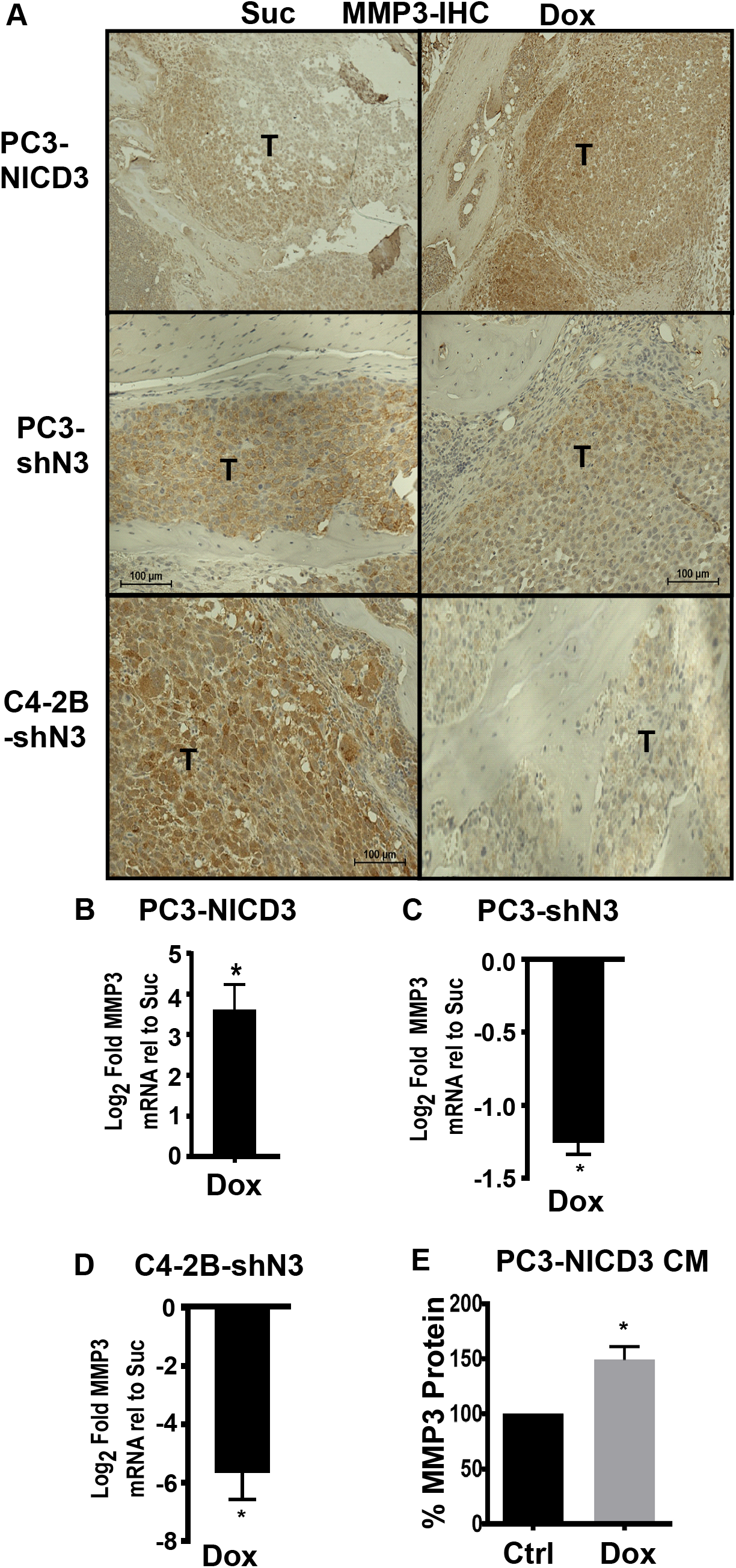
Notch3 promotes the expression of MMP-3. **(A)** Tibiae from mice injected with PC3-NICD3, PC3-shN3, or C4-2B-shN3 cells treated with sucrose (Suc) or doxycycline (Dox) were stained with human-specific MMP-3 antibody. T=tumor **(B-D)** Levels of human-specific MMP-3 mRNA from tibiae in (A) assessed by qRT-PCR. Expressed Log_2_ fold relative to sucrose controls. **(E)** Relative levels of MMP-3 in conditioned medium (CM) from PC3-NICD3 cells treated with vehicle (Ctrl) or doxycycline (Dox) assessed by ELISA. Error bars are S.E.M, n>4; *0.01≤p≤0.05.

### Notch3, in cancer cells, induces osteoblasts and inhibits osteoclasts

Two ways in which NICD3, acting in tumor cells, could inhibit osteolytic lesions is through a decrease in osteoclastogenesis or through an increase in osteoblastogenesis. To evaluate these possibilities, we analyzed expression of osteoclast inhibitors in PC3-NICD3 tumor-bearing tibiae. Mouse-specific IL-10 mRNA, an inhibitor of osteoclastogenesis (26, 27), is upregulated over 8-fold in Dox-treated PC3-NICD3 tumors relative to sucrose-treated controls (Fig. 4A). In addition, the ratio of mouse-specific OPG/RANKL mRNA, which limits osteoclastogenesis (28), is elevated over 2-fold in PC3-NICD3 tibiae (Fig. 4B). This increase in osteoclastogenesis inhibitors by NICD3 *in vivo* supports the observed decrease in active TRAP-positive osteoclasts in the NICD3-expressing tumors (see Fig. 1F). Conversely, there was a significant 5-8-fold increase in osteoblast markers, bone sialoprotein (BSP), osteocalcin (OCN), and alkaline phosphatase (ALP) (Fig 4C), in the Dox-treated PC3-NICD3 tumors. This increase in osteoblast markers by NICD3 *in vivo* supports the observed increase in osteoblasts in the NICD3-expressing tumors (see Fig. 1G). Furthermore, OCN is a marker of mature osteoblasts, and its induction by NICD3 indicates NICD3 promotes osteoblast differentiation *in vivo.* Thus, NICD3 expression in tumors leads to increased osteoblastogenesis and decreased osteoclastogenesis *in vivo.*

**Figure 4.**
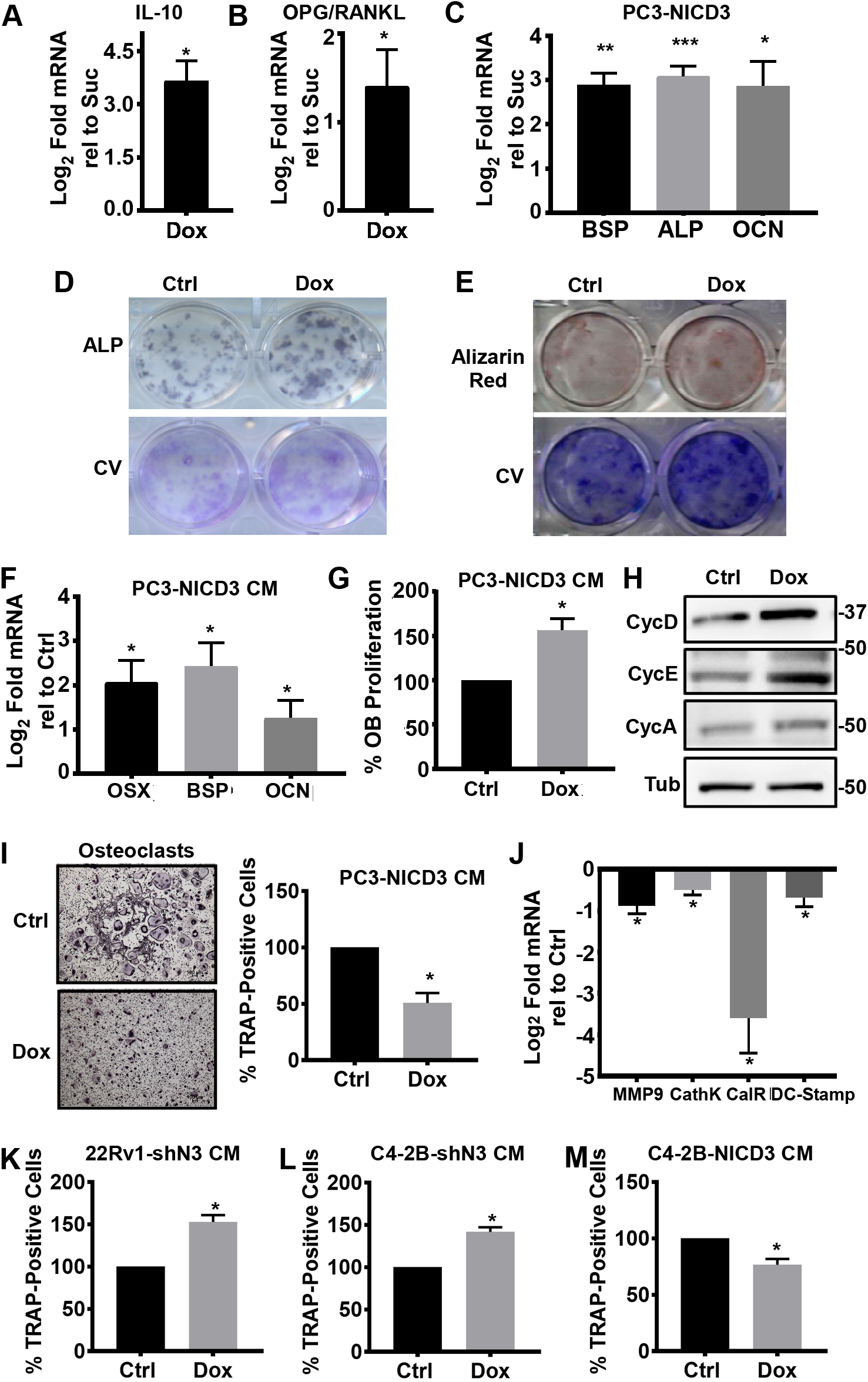
NICD3 inhibits osteoclastogenesis and promotes osteoblastogenesis. **(A-C)** Expression of mouse-specific mRNA isolated *in vivo* from tibiae of mice injected with PC3-NICD3 cells and treated with sucrose (Suc) or doxycycline (Dox): **(A)** IL-10, **(B)** OPG/RANKL ratio, and **(C)** bone sialoprotein (BSP), alkaline phosphatase (ALP), or osteocalcin (OCN). Expressed Log_2_ fold relative to sucrose controls. **(D-H)** Bone marrow-derived osteoblasts differentiated in the presence of conditioned medium (CM) from PC3-NICD3 cells treated with doxycycline (Dox) or vehicle (Ctrl). **(D)** Colonies were stained for ALP or crystal violet (CV). **(E)** Colonies were stained for Alizarin red or crystal violet (CV). **(F)** Expression of mouse-specific mRNA from treated osteoblast cultures: osterix (OX), bone sialoprotein (BSP), or osteocalcin (OCN). Expressed Log_2_ fold relative to vehicle controls. **(G)** MTT assay of treated osteoblast cultures. **(H)** Levels of Cyclin A, D, E and tubulin (Tub) from treated osteoblast cultures assessed by immunoblotting. **(I-J)** Bone marrow-derived osteoclasts differentiated in the presence of conditioned medium (CM) from doxycycline (Dox) or vehicle-treated (Ctrl) PC3-NICD3 cells. **(I)** TRAP+ cells with 2 > nuclei quantified. **(J)** Expression of mouse-specific mRNA from osteoclast cultures: MMP9, cathepsin K (CathK), calcitonin receptor (CR), or DC-Stamp. **(K-M)** Bone marrow-derived osteoclasts differentiated in the presence of conditioned medium (CM) from **(K)** 22Rv1-shN3 (Notch3 shRNA), **(L)** C4-2B-shN3, and **(M)** C4-2B-NICD3 expressing cells. Percentage of TRAP+ cells quantified. Error bars are S.E.M, n=3; *0.01≤p≤0.05; **0.001≤p<0.01; ***p<0.001.

To more directly measure the effect of the NICD3-expressing tumor cells on osteoblasts, conditioned medium (CM) from Dox-treated PC3-NICD3 cells in culture was added to differentiating osteoblasts *in vitro.* CM from Dox-stimulated PC3-NICD3 cells enhanced osteoblast differentiation as measured by increased alkaline phosphatase (ALP) staining, increased colony formation (Fig. 4D), and mineralization as detected by Alizarin red staining (Fig 4E). This was accompanied by a 3-4-fold increase in osteoblast differentiation markers, i.e. Osterix (OSX), BSP, and OCN (Fig. 4F). CM from Dox-treated PC3-NICD3 cells also stimulated osteoblast proliferation (Fig. 4G) and expression of Cyclin A, D and E (Fig. 4H).

To investigate how Notch3 expression in cancer cells affects osteoclasts, we added CM from Dox-treated PC3-NICD3 cells to differentiating osteoclasts. NICD3 CM inhibited the formation of active osteoclasts as measured by the reduced number of fused multinucleated TRAP-positive cells (Fig. 4I) and reduced mRNA expression of late osteoclast differentiation markers, particularly the calcitonin receptor (Fig. 4J). Reciprocally, CM from Dox-treated C4-2B-shN3 or 22Rv1-shN3 cells stimulated the formation of TRAP-positive multinucleate osteoclasts (Fig. 4K-L). Finally, CM from Dox-treated C4-2B-NICD3 cells decreased the number of TRAP-positive multinucleate osteoclasts (Fig. 4M). We did not observe any effects of the CM on osteoclast survival; CM did not induce caspase3 cleavage as assessed by immunostaining (not shown). Together these results indicate that CM from Notch3-expressing cells inhibits osteoclast differentiation while simultaneously stimulating osteoblast proliferation and differentiation. These effects may explain the decrease in osteolytic lesion formation seen in Notch3-expressing tumors.

### Notch3 inhibits osteolytic bone lesion development in an MMP-3-dependent manner

To determine if Notch3 induction of MMP-3 is responsible for the observed effects of NICD3 conditioned medium (CM) on osteoblasts and osteoclasts, we directly tested the ability of human recombinant MMP-3 (rMMP3) to regulate differentiation. The addition of rMMP3 to differentiating osteoclasts reduced the formation of TRAP-positive multinucleate cells (Fig. 5A), but had no effect on osteoblast differentiation as measured by ALP staining (Fig. 5B). However, rMMP3 did promote osteoblast proliferation (Fig. 5C).

**Figure 5.**
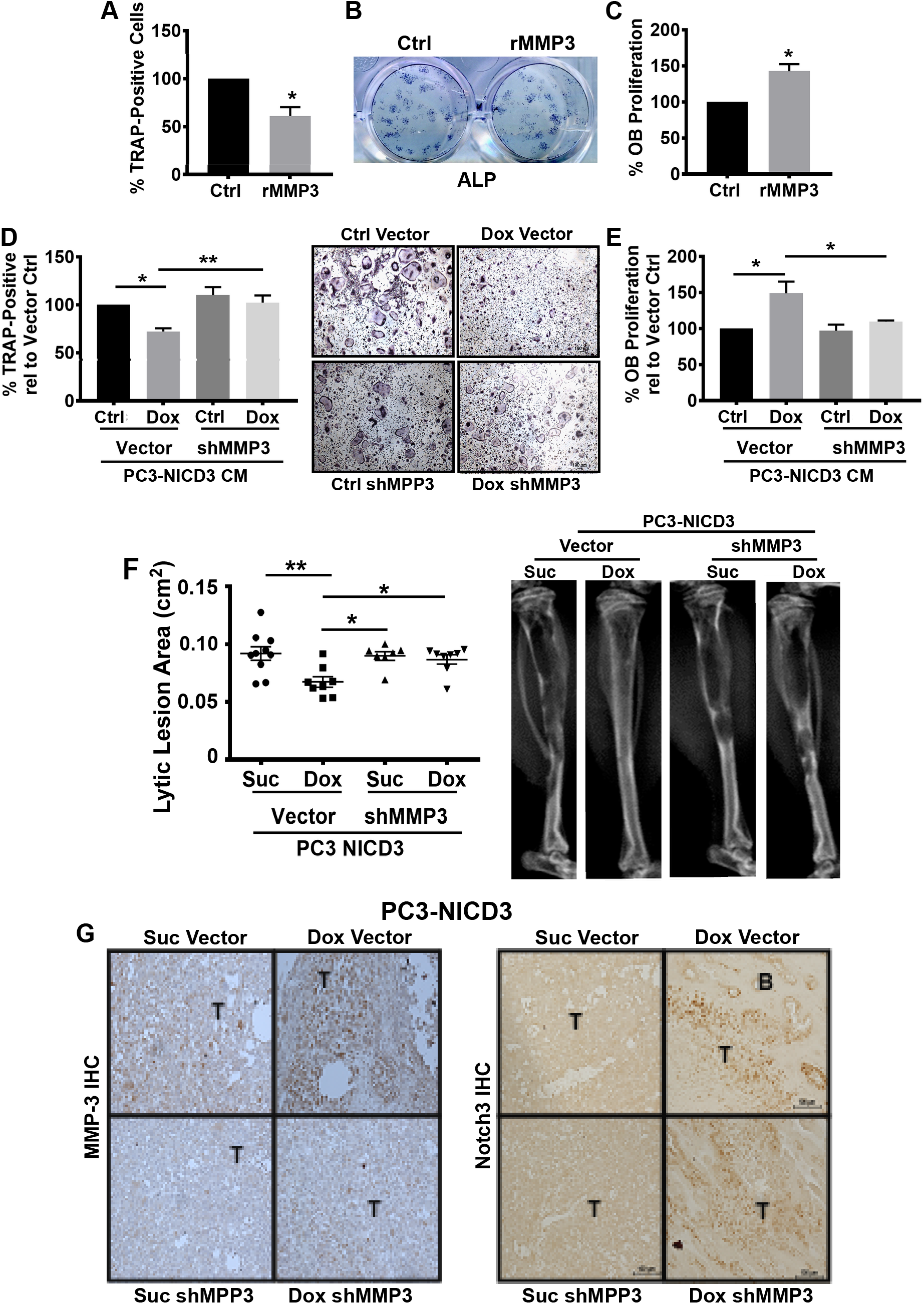
Notch3 promotes osteoblastic lesion development in an MMP-3-dependent manner. **(A)** Bone marrow-derived osteoclasts differentiated in the presence of 25 ng/ml recombinant human MMP-3 (rMMP3). Percentage of TRAP+ cells with 2 > nuclei quantified. **(B-C)** Bone marrow-derived osteoblasts differentiated in the presence of recombinant MMP-3 (rMMP3) and **(B)** immunostained for ALP or **(C)** proliferation measured by MTT assay. **(D-F)** Tet-inducible PC3-NICD3 cells engineered to stably express MMP-3 shRNA (PC3-NICD3-shMMP3). **(D)** Bone marrow-derived osteoclasts or **(E)** osteoblasts differentiated in the presence of conditioned medium (CM) from doxycycline-treated (Dox) or vehicle-treated (Ctrl) PC3-NICD3-vector or PC3-NICD3-shMMP3 cells. **(D)** Percentage of TRAP+ cells quantified. **(E)** MTT assay of treated osteoblasts. **(F)** Tet-inducible PC3-NICD3-vector and PC3-NICD3-shMMP3 injected into tibiae of mice treated with doxycycline (Dox) or sucrose (Suc). Lytic lesion area on X-ray quantified. **(G)** Tibial tumors from **(F)** immunostained (IHC) with Notch3 or MMP-3 antibodies. Error bars are S.E.M, n>8; *0.01≥ p≤0.05; **0.001≤p<0.01.

To investigate the dependency of NICD3 on MMP-3 for its inhibitory effects on osteoclasts, PC3-NICD3 cells were engineered to stably express constitutive MMP-3 shRNA (shMMP3). CM from Dox-treated PC3-NICD3-shMMP3 cells rescued the block in TRAP-positive multinucleate cell differentiation induced by CM from Dox-treated PC3-NICD3-vector cells (Fig. 5D). Addition of rMMP3 restored the decrease in osteoclast differentiation that was blocked by shMMP3 (Supplementary Figure S6A), indicating MMP-3 is responsible for decreasing osteoclast differentiation. CM from Dox-treated PC3-NICD3-shMMP3 cells also prevented the promotion of osteoblast proliferation (Fig. 5E) and this was partially reversed by adding rMMP3 (Supplementary Figure S6B). We tested whether shMMP3 could also rescue osteolytic lesion formation. Dox-treated PC3-NICD3-shMMP3 tibiae had increased osteolytic lesion area relative to Dox-treated PC3-NICD3-vector (Fig. 5F). MMP-3 knock-down and NICD3 induction was validated by IHC staining of tibiae (Fig. 5G). While tumor proliferation was decreased ~20% in the PC3-NICD3-shMMP3 tumors relative to PC3-NICD3-vector tumors, this was independent of NICD3 expression (Supplementary Fig. S3E). Loss of MMP-3 did not alter the proliferation of cells grown *in vitro* (Supplementary Figure S4D).

Thus, rescue of the osteolytic phenotype was not due to enhanced tumor or tumor cell proliferation. Altogether, these data indicate MMP-3 is required for Notch3-mediated inhibition of osteolytic bone lesion development by prostate tumors, where it acts primarily through inhibition of osteoclast differentiation and contributes to osteoblast proliferation.

### Notch3 and MMP-3 expression is elevated in human bone metastases

Notch signaling pathway components are upregulated in Gleason 8 compared to Gleason 6 tumors (17). Furthermore, expression of Notch3 in prostate tumors is inversely correlated with patient survival (29), indicating the likely importance of Notch3 in prostate cancer progression. To measure the levels of Notch3 and MMP-3 in prostate cancer bone metastases, two human tissue microarrays, TMA-170 with 10 matched visceral and bone metastases, and UWTMA79 with 30 matched visceral and bone metastases, were analyzed by IHC with antibodies to Notch3 or MMP-3. Bone metastases in both MTAs expressed more Notch3 and MMP-3 compared to matched visceral metastases (Fig. 6A,B). Furthermore, there was a significant correlation between expression of Notch3 and MMP-3 in all metastases (Fig. 6C).

**Figure 6.**
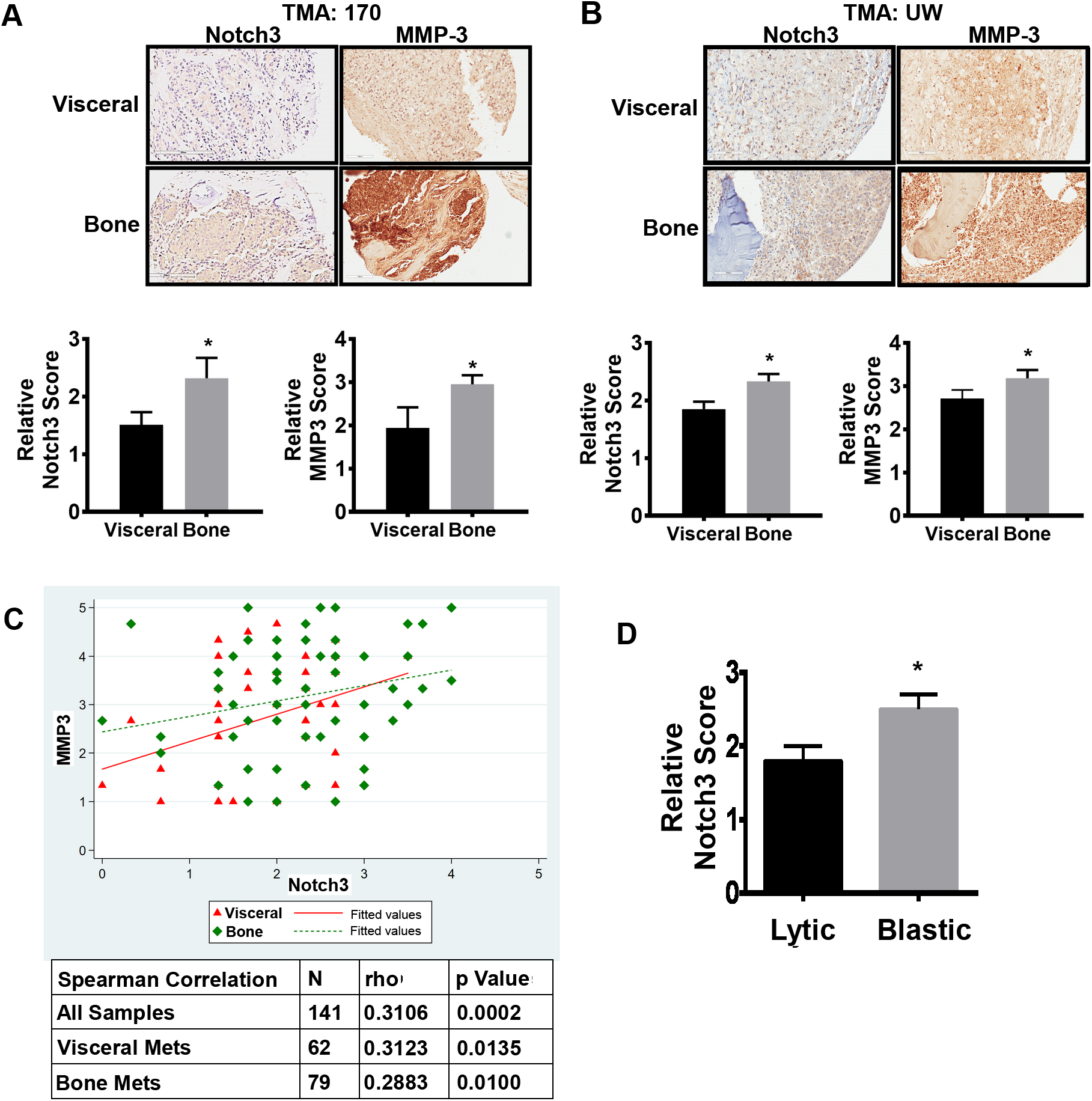
Notch3 and MMP-3 expression is elevated in bone metastases. Tissue microarrays (A) TMA-170 and (B) UWTMA79 probed for Notch3 and MMP-3 expression by IHC. Levels of expression were compared between patient-matched visceral and bone metastases. (C) Frequency of Notch3 and MMP-3 co-expression across all samples. (D) UWTMA79 bone metastases were assessed for lytic (n=18) versus osteoblastic (n=42) lesions. Association of elevated Notch3 or MMP-3 levels with lesion type was assessed. Error bars are S.E.M; *0.01≤ p≤0.05.

We also found that high Notch3 expression was more highly associated with osteoblastic lesions compared to lytic lesions (Fig. 6D). Altogether, our findings indicate that a Notch3-MMP-3 axis plays an integral part in favoring osteoblastic lesion formation in prostate cancer bone metastasis.

## DISCUSSION

Our studies demonstrate that Notch3 activation in prostate cancer bone metastases suppresses osteolytic lesions through induction and secretion of MMP-3, which acts to suppress osteoclast differentiation and enhance osteoblast proliferation. Notch3 expression increases with Gleason grade, and is associated with prostate cancer recurrence, suggesting its strong importance in cancer progression (16, 18). This is the first study to demonstrate that Notch3 is highly expressed in human prostate cancer-induced bone metastasis as compared to visceral metastasis and preferentially in osteoblastic bone metastases, relative to osteolytic lesions. Interestingly, Notch3 is activated by hypoxia, and bone metastases are highly hypoxic (18, 30). The hypoxic bone environment could be a driver of elevated Notch3 expression.

The promotion of osteoblast proliferation and differentiation by simply inducing NICD3 expression in an osteolytic cell line, suggests Notch3 is a key regulator of the osteoblastic phenotype that is so unique to prostate cancer. While prostate cancer is highly osteoblastic, there are also underlying osteolytic events (4, 5, 31). Our findings indicate that elevated Notch3 reduces the osteolytic events through its extrinsic effects on the bone microenvironment, with little to no intrinsic effect on tumor cell proliferation. NICD3 accomplishes this by promoting osteoblast proliferation and differentiation, while simultaneously inhibiting osteoclastogenesis; thus, tipping bone remodeling towards enhanced bone formation. We further found that this osteolytic suppression is dependent on MMP-3. The ability of NICD3 to suppress osteoclasts and induce osteoblast proliferation, but not osteoblast differentiation, was mediated by MMP-3. Thus, Notch3 is likely operating on the bone tumor microenvironment through at least one other mechanism.

MMP-3 is known to promote the cleavage of PTHrP produced by cancer cells, which stimulates osteoblast migration and mineralization *in vitro* but lacks the osteolytic function associated with full length PTHrP (32). PTHrP also promotes osteoblast proliferation (33). Thus, the effects of the Notch3-MMP-3 axis on osteoblasts might depend in part on MMP-3-cleaved PTHrP. A previous study demonstrated that MMP-3 induces osteoclast death (34); however, we did not detect induction of any caspase-dependent cell death in our in vitro cultures either from NICD3-conditioned medium or by recombinant MMP-3.

MMP-3 is highly expressed in other cancers like breast, lung, pancreas, and osteosarcoma, which correlates with poor patient survival (35, 36). In prostate cancer, MMP-3 expression is higher in primary tumors compared to PIN and normal tissue (37), and MMP-3 was previously reported to be elevated in a prostate cancer bone xenograft, but the source or function was not identified (38). MMP-3 levels are reportedly higher in the serum of prostate cancer patients with bone metastatic disease (24). We are the first to demonstrate both elevated MMP-3 and Notch3 expression in human bone metastases, compared to visceral metastases, in prostate cancer patients. Interestingly, MMP-3 expression is induced in osteoblasts in response to inflammatory signals or mechanical stress and is necessary for bone remodeling (39, 40) and MMP-3 is elevated in the serum of rheumatoid arthritis patients, which is accompanied by both bone remodeling and inflammation (41). Thus, tumors that secret MMP-3 may access this normal bone stress response. This is the first study to demonstrate how MMP-3, secreted from cancer cells, modulates the function of bone cells in prostate cancer bone lesions.

In the past ten years there have been improvements in the options to treat bone metastatic prostate cancer; however, the available drugs (docetaxel, carbitaxel, abiraterone or enzalutamide) only increase patient survival for a short time as most patients eventually develop resistance to these drugs and finally succumb to the disease (1, 42). Our data demonstrates that NICD3 modulates the bone microenvironment, but has minimal effects on tumor cell proliferation. This suggests that inhibitors of Notch alone might not be effective at reducing tumor burden. Furthermore, while inhibition of Notch3 activity will decrease osteoblastic lesions, it may enhance lytic lesions by promoting osteoclast function and inhibiting the osteoblasts. Increasing the function of osteoclasts in osteoblastic metastasis could be an important step towards rebalancing bone homeostasis; however, there will be a need for combinatorial therapy to inhibit the excessive osteolysis caused by Notch inhibition.

New γ-secretase inhibitors (pan-Notch inhibitors) in combination with bicalutamide have been in clinical trials but were terminated with no useful information (43). Recently, Cui et al demonstrated that a γ-secretase inhibitor in combination with docetaxel showed greater antitumor activity than either agent alone in DU145 prostate cancer tibial xenografts (20). Thus, to effectively reduce both tumor growth and block abnormal bone remodeling, it will be necessary to use combination treatments that target both the tumor and the tumor microenvironment.

## MATERIALS AND METHODS

### Cells lines and animals

PC3 and 22Rv1 cells were obtained from ATCC, and C4-2B cells was obtained from Dr. Robert Sikes (University of Delaware) (44). All lines were maintained in RPMI1640 (Gibco) supplemented with 10%FBS (Gemini). All lines were validated by STR analysis yearly and checked for mycoplasma contamination using MycoAlert PLUS kit (Lonza). Conditioned medium was collected by starving sub-confluent cells for 24 h in α-MEM (Gibco).

The 6-week old male NSG mice used in this study were bred and maintained in a pathogen-free and ALAAC-certified barrier facility following guidelines from VARI Institutional Animal Care and Use Committee and the Department of Defense (DOD) Animal Care and Use Committee.

### Doxycycline-inducible shRNA and NICD

Doxycycline-inducible shRNA targeting Notch3 was generated as previously described (14, 45). NICD1 in EF.hICN1.CMV.GFP (gift from Linzhao Chen (46); Addgene #17623) was PCR amplified with Syzygy SyFi high fidelity polymerase (Integrated Scientific Solutions, SY-100) using the following primers: Fwd 5’-AATTGTCGACCCAAGCTGGCTAGTTAAGC-3’ and Rev 5’-TTAAGCGGCCGCTTTATTCCAGCACACTGGCGGC-3’. PCR product was ligated with LigateIT rapid ligase (Affymetrix) into pENTR3C (Invitrogen) between SalI and NotI sites, creating pENTR3C-NICD1. NICD3 was similarly PCR subcloned from pCLE-NICD3 (gift from Nicholas Gaiano (47); Addgene #26894) into pENTR3C between SalI and NotI, creating pENTR3C-NICD3, using primers: Fwd 5’-AATTGTCGACCCCGCCTCTAGCACTTTG G-3’ and Rev 5’-TTAAGCGGCCGCTTTATTCGATCTAAGAACTGACGAGCG-3’.

pENTR3C-NICD1 and pENTR3C-NICD3 were recombined via L/R Clonase II (Thermo) into pLenti-CMV-Puro-DEST (w118-1, gift from Eric Campeau and Paul Kaufman (48); Addgene #17452) to make pLenti-Tet-NICD1-Puro and pLenti-Tet-NICD3-Puro. Integrity of the plasmids verified by sequencing.

PC3 cells were first infected with pLenti-CMV-TetR-Blast (716-1, gift from Eric Campeau and Paul Kaufmann (48); Addgene #17492) and a stable pooled TetR line was created by selecting in 2 μg/ml blasticidin. TetR line was then infected with NICD1 or NICD3-expressing lentivirus and pooled clones selected in 2 μg/ml puromycin. Cell lines targeting MMP-3 with shRNA (shMMP3) were generated by purchasing a lentiviral shRNA knockdown vector containing shMMP3 sequence (pLV[shRNA]-Hygro-U6>hMMP3) from Vector Builder. Targeting sequence was 5’-AGAGTAACAGCTGGCTTAATT-3’. Pooled cell lines were selected in 100 μg/ml hygromycin. Cell lines were treated with increasing concentrations of Doxycycline (Millipore) to test for optimal shNotch3, NICD1, or NICD3 induction by immunoblotting. MTT assay were performed on PC3-NICD3 vector versus PC3-NIC3-shMMP3 cells or after treating PC3-NICD3 cells with or without 500ng/ml of Doxycycline for 1d, and 22Rv1 or C4-2B shNotch3 cells with or without 10ng/ml of Doxycycline after 7 days in culture, in triplicate using the manufacturers protocol (ATCC).

### Mouse injections and radiographic imaging

One million PC3 or C4-2B cells or 0.2 x 10^6^ 22Rv1 cells resuspended in 10 μl of PBS were injected into the tibiae of 6-week old male NSG mice. As a control, 10 μl of PBS was injected into the contralateral tibia. Each experiment had a minimum of 7, and up to 18, mice per cohort. The number of mice per cohort was based on the known *in vivo* growth rate of the different cell lines and sample power was not predetermined. Half the mice injected with shNotch3, NICD1, or NICD3 expressing cells were non-blindly randomly assigned to receive 1 mg/ml Doxycycline in the drinking water supplemented with 5% Sucrose starting from the 1st day of injection and changed weekly. Control mice received only 5% sucrose. Mice were blindly radiographically imaged weekly using Bioptics piXarray Digital Specimen Radiography (Faxitron Bioptics). The lytic bone lesions were scored in a blinded manner by measuring the area of all visible lesions using MetaMorph quantitative image analysis software (Molecular Devices, Inc.). At the end-point, ~3-4 weeks for PC3, ~8 weeks for C4-2B and 22Rv1, tibiae were snap frozen for RNA and protein analysis or fixed in formalin for immunohistochemistry. No mice were excluded from the analyses.

For subcutaneous injections, 1.0 x 10^6^ PC3, C4-2B, or 22Rv1 cells resuspended in 100 μl of PBS were injected subcutaneously into the flank area of 6-week old male NSG mice and tumors harvested after 4 or 8 weeks and frozen in liquid nitrogen.

### Bone marrow differentiation, mineralization, and proliferation assays

#### Osteoblasts

Mouse bone marrow cells extracted from the long bones of 5-6 week old naïve NSG mice, were plated in α-MEM medium supplemented with 10%FBS, left to adhere for 3 d, and then differentiated into osteoblasts using 50 μg/ml Vitamin C for another 7 d for ALP staining and 14 d for Alizarin red staining. The medium was changed every 3 d. On the third day of culture, bone marrow cells were treated with conditioned medium from prostate cancer cell lines bearing Tet-inducible shNotch3 or NICD3 treated with or without Dox. On the last 3 days, cells were serum-starved. Alkaline phosphatase (ALP) staining were performed by fixing cells in 10% neutral buffered formalin and then staining with NBT/BCIP Substrate Solution (Thermo Scientific) at 37^0^C. Alizarin red staining was performed by fixing cells in 10% neutral buffered formalin and then staining with Alizarin red solution (Millipore) at room temperature. Osteoblast colony formation was monitored by staining duplicate wells with 0.5% crystal violet. Osteoblast proliferation was evaluated via MTT assay in triplicate using the manufacturers protocol (ATCC).

#### Osteoclasts

Bone marrow cells were plated in α-MEM supplemented with 10% FBS. After 24 hours, unattached cells were replated and cultured in α-MEM supplemented with 10% FBS and 25 ng/ml macrophage colony-stimulating factor (mCSF). Two days later, cells were supplemented with 25 ng/ml mCSF and 30 ng/ml RANKL (R&D) for another 4 d. During these last 4 days, cells were treated with conditioned medium from cell lines bearing Tet-inducible shNotch3 or NICD3 treated with or without Dox. Cells were serum-starved for the last 24 h of culture. TRAP staining was performed according to manufactures protocol (Sigma Aldrich). TRAP-positive multinucleate (2 ≥ nuclei/cell) osteoclasts were counted in 10 random fields. In a subset of assays, cultured osteoblasts or osteoclasts were treated with 25 ng/ml recombinant full-length human MMP-3 (Abcam, ab96555) in place of conditioned medium or in combination with conditioned medium.

### Histology and immunohistochemistry

Mouse tibiae were fixed in 10% neutral-buffered formalin (Sigma) for 4 d at 4°C, followed by decalcification in 14% EDTA for 5-6 d at 4°C, and then embedded in paraffin. Serial bone sections 5 μm thick were used for lHC staining. Hematoxylin and eosin and TRAP staining were done as previously reported (49). Paraffin embedded sections were stained using antibodies against Ki67 (1:100; Thermo Scientific; RM-9106), MMP-3 (1:100; Abcam; ab52915), or Notch3 (1:300; Santa Cruz; sc5593).

### TRAP, Osteoblasts, and Ki67 quantification

Serial sections stained for TRAP were scanned and images collected using Aperio software. TRAP-positive multinucleate osteoclasts (2 ≥ nuclei/cell) at the bone/tumor interface were counted after scanning slides in Aperio and counted at 20X magnification using Imagescope software. H&E counter-stained sections from TRAP stained sections were used to count osteoblasts. Osteoblasts were identified by their distinct morphology along the bone-tumor interface. Ki67 was quantified blinded, by counting 3-4 random fields under 400X magnification.

### TMA staining and quantification

Sections of a 45-patient tissue microarray (TMA) having 30 patient-matched visceral and bone metastases was obtained from the Prostate Cancer Biorepository Network (PCBN), UWTMA79. The TMA-170 microarray of 12 patients, consisting of 10 patient-matched visceral and bone metastases, was prepared from tissues obtained from the rapid autopsy program at University of Michigan. Both TMAs were stained using MMP-3 (1:75, Abcam; ab52915) and Notch3 (1:1000; Protein Tech 55114-1-AP) antibodies.

Two board-certified pathologists, without any prior knowledge of the patients’ clinical information, independently and blindly used H and modified H scoring to quantify the TMAs for MMP-3 and Notch3 expression in triplicate. Average scores were calculated for each tumor and pathologist. Pathological analysis of the PCBN array identified lytic, predominantly lytic, predominantly blastic, blastic, mixed, and penic/porotic lesions.

### Protein extraction, immunoblot, ELISA, and cytokine array

Tumor-containing tibiae and subcutaneous tumors were snap-frozen in liquid nitrogen. Frozen tibiae were homogenized in RIPA buffer (50) using a FastPrep-24 tissue homogenizer (MP Biomedicals). Prostate cancer cell lines were lysed using RIPA buffer as described previously (50). Total protein (10-20 μg) was separated by SDS-PAGE and transferred to PVDF membranes (Fisher). Membranes were blocked with 5% BSA-TBST and incubated with antibodies diluted in 5% BSA-TBST as previously described (50). Primary antibodies were detected using HRP-conjugated secondary antibodies (Sigma) by chemiluminescence using Quantity One imaging software on a Bio-Rad Gel Docking system. Antibodies listed in Supplementary Table S1.

Equal amounts of protein from subcutaneous or tibiae tumors pooled from two mice were incubated with a 174-protein spotted human cytokine antibody array C2000 (Ray Biotech Inc) according to the manufacturer’s protocol.

An MMP-3 ELISA (R&D) that recognizes both the proenzyme and active form was performed on 24 h serum-starved PC3-NICD3 cells treated with or without Dox. Media were collected and concentrated (Ultracel-10k; Millipore) and equal volumes were used to quantify total MMP-3.

### RNA extraction and qRT-PCR

Total RNA was extracted from frozen tumor-containing tibiae or cell lines using TRIzol (Invitrogen, Carlsbad, CA). Equal amounts of RNA were reverse transcribed through the SuperScript first-strand synthesis system (Invitrogen) and qRT-PCR was performed using SYBR Green Supermix (Bio-Rad, Hercules, CA) on Applied BioSystems. Primers were synthesized by Integrated DNA Technologies (IDT) and the levels of target mRNAs were standardized to GAPDH and plotted as Log_2-_fold. Primers listed in Supplementary Table S2.

### Statistical analysis

All statistical analyses on the mice with one or two comparison groups (Fig 1–4) used Student’s t-test and in experiments where more than two groups (Fig. 5) were being compared, ANOVA was used. In all analyses, two-tailed significance is reported. For statistical analysis of the TMAs, multivariate analysis of Notch3 and MMP-3 expression levels in visceral versus bone metastases was conducted using mixed effect models. The mixed effect models account for the correlation between the measurements obtained across multiple observations per individuals. Spearman correlation was used to calculate correlation between MMP-3 and Notch3 expression across all metastases. A regression model controlling for repeated measures by patient was fit to test the difference in Notch3 or MMP-3 expression between osteolytic and osteoblast groups of bone metastases. For this analysis, the lytic and predominantly lytic (n=18 lytic) and the blastic and predominantly blastic (n=42 blastic) were pooled into 2 groups. The mixed (n=3) and penic/porotic (n=12) lesions were excluded from the analysis. All analyses were conducted using STATA 15 and met the expected distribution. The variance between groups for both the mice and tissue studies was similar.

## ACKNOWLEDGEMENTS

Funding for this project was provided by a DOD Postdoctoral Fellowship W81XWH-16-1-0136 (SSG), DOD W81XWH-14-1-0479 (SBF, LT, CKM), University of Arizona Cancer Center (UACC) NIH/NCI P30CA023074 (KS, CKM), the Van Andel Research Institute (VARI) (GH, XL), and the University of Arizona (CKM). The TMA obtained from the Prostate Cancer Biorepository Network (PCBN) was supported by funds from the DOD Prostate Cancer Research Program: W81XWH-14-2-0182, W81XWH-14-2-0183, W81XWH-14-2-0185, W81XWH-14-2-0186, and W81XWH-15-2-0062. We wish to thank the following: Dr. Denise Roe, Director of Biostatistics Shared Resource at UACC for biostatistics support; Dr. Chunyan Liu at Ventana Medical Services, Oro Valley, AZ for her pathology services; Dr. Colm Morrissey in the Department of Urology, University of Washington, Seattle WA, for his assistance with providing pathological assessment of bone metastasis lesions in the PCBN TMA. Alexandra VanderArk, Veronique Schulz, and Ghada Y.T Mohsen at Van Andel Research Institute for their technical expertise; Zachary Madaj for statistical analysis expertise; Lisa Turner, Kristin Feenstra and Bree Berghuis of the VARI Pathology and Biorepository Core for their pathology and Aperio expertise; Su Yanli and Staff of the VARI Vivarium and Transgenics core for technical assistance with animal experiments; David Nadziejka for technical editing of the manuscript; Jeanie Wedberg and Michelle Minard at the Van Andel Institute and David Alvarado at University of Arizona for their administrative support.

## REFERENCES

1. Ganguly SS, Li X, Miranti CK. The host microenvironment influences prostate cancer invasion, systemic spread, bone colonization, and osteoblastic metastasis. Front Oncol. 2014;4:364.

2. Siegel RL MK, Jemal A. Cancer statistics, 2015. CA Cancer J Clin. 2015;65(1):5–29.

3. Karayi MK, Markham AF. Molecular biology of prostate cancer. Prostate Cancer Prostatic Dis. 2004;7(1):6–20.

4. Roudier MP, Morrissey C, True LD, Higano CS, Vessella RL, Ott SM. Histopathological assessment of prostate cancer bone osteoblastic metastases. J Urol. 2008;180(3):1154–60.

5. Sottnik JL, Keller ET. Understanding and targeting osteoclastic activity in prostate cancer bone metastases. Curr Mol Med. 2013;13(4):626–39.

6. Jin JK, Dayyani F, Gallick GE. Steps in prostate cancer progression that lead to bone metastasis. Int J Cancer. 2011;128(11):2545–61.

7. Frank SB, Miranti CK. Disruption of prostate epithelial differentiation pathways and prostate cancer development. Front Oncol. 2013;3:273.

8. Kopan R, Ilagan MX. The canonical Notch signaling pathway: unfolding the activation mechanism. Cell. 2009;137(2):216–33.

9. Zanotti S, Canalis E. Notch and the skeleton. Mol Cell Biol. 2010;30(4):886–96.

10. Wang XD, Shou J, Wong P, French DM, Gao WQ. Notch1-expressing cells are indispensable for prostatic branching morphogenesis during development and re-growth following castration and androgen replacement. J Biol Chem. 2004;279(23):24733–44.

11. Kwon OJ, Valdez JM, Zhang L, Zhang B, Wei X, Su Q, et al. Increased Notch signalling inhibits anoikis and stimulates proliferation of prostate luminal epithelial cells. Nature communications. 2014;5:4416.

12. Bin Hafeez B, Adhami VM, Asim M, Siddiqui IA, Bhat KM, Zhong W, et al. Targeted knockdown of Notch1 inhibits invasion of human prostate cancer cells concomitant with inhibition of matrix metalloproteinase-9 and urokinase plasminogen activator. Clin Cancer Res. 2009;15(2):452–9.

13. Ye QF, Zhang YC, Peng XQ, Long Z, Ming YZ, He LY. Silencing Notch-1 induces apoptosis and increases the chemosensitivity of prostate cancer cells to docetaxel through Bcl-2 and Bax. Oncol Lett. 2012;3(4):879–84.

14. Frank SB, Berger PL, Ljungman M, Miranti CK. Human prostate luminal cell differentiation requires NOTCH3 induction by p38-MAPK and MYC. J Cell Sci. 2017;130(11):1952–64.

15. Zhang S, Chung WC, Wu G, Egan SE, Xu K. Tumor-suppressive activity of Lunatic Fringe in prostate through differential modulation of Notch receptor activation. Neoplasia. 2014;16(2):158–67.

16. Long Q, Johnson BA, Osunkoya AO, Lai YH, Zhou W, Abramovitz M, et al. Protein-coding and microRNA biomarkers of recurrence of prostate cancer following radical prostatectomy. Am J Pathol. 2011;179(1):46–54.

17. Ross AE, Marchionni L, Vuica-Ross M, Cheadle C, Fan J, Berman DM, et al. Gene expression pathways of high grade localized prostate cancer. Prostate. 2011;71(14):1568–77.

18. Danza G, Di Serio C, Ambrosio MR, Sturli N, Lonetto G, Rosati F, et al. Notch3 is activated by chronic hypoxia and contributes to the progression of human prostate cancer. Int J Cancer. 2013;133(11):2577–86.

19. Cui J, Wang Y, Dong B, Qin L, Wang C, Zhou P, et al. Pharmacological inhibition of the Notch pathway enhances the efficacy of androgen deprivation therapy for prostate cancer. Int J Cancer. 2018.

20. Cui D, Dai J, Keller JM, Mizokami A, Xia S, Keller ET. Notch pathway inhibition using PF-03084014, a gamma-secretase inhibitor (GSI), enhances the antitumor effect of docetaxel in prostate cancer. Clin Cancer Res. 2015;21(20):4619–29.

21. Zhang Z, Wang H, Ikeda S, Fahey F, Bielenberg D, Smits P, et al. Notch3 in human breast cancer cell lines regulates osteoblast-cancer cell interactions and osteolytic bone metastasis. Am J Pathol. 2010;177(3):1459–69.

22. Sethi N, Dai X, Winter CG, Kang Y. Tumor-derived JAGGED1 promotes osteolytic bone metastasis of breast cancer by engaging notch signaling in bone cells. Cancer Cell. 2011;19(2):192–205.

23. Gong Y, Chippada-Venkata UD, Oh WK. Roles of matrix metalloproteinases and their natural inhibitors in prostate cancer progression. Cancers (Basel). 2014;6(3):1298–327.

24. Srivastava P, Kapoor R, Mittal RD. Impact of MMP-3 and TIMP-3 gene polymorphisms on prostate cancer susceptibility in North Indian cohort. Gene. 2013;530(2):273–7.

25. Jung K. NL, Lein M., Priem F., Schnorr D., Loening S.A. Matrix metalloproteinases 1 and 3, tissue inhibitor of metalloproteinase-1 and the complex of metalloproteinase-1/tissue inhibitor in plasma of patients with prostate cancer. Int J Cancer. 1997;74:220–3.

26. Hong MH WH, Jin CH, Pike jw. The inhibitory effect of interleukin-10 on mouse osteoclast formation involves novel tyrosine-phosphorylated proteins. J Bone Miner Res. 2000;15(5):911–8.

27. Evans KE, Fox SW. Interleukin-10 inhibits osteoclastogenesis by reducing NFATc1 expression and preventing its translocation to the nucleus. BMC Cell Biol. 2007;8:4.

28. Boyce BF, Xing L. Biology of RANK, RANKL, and osteoprotegerin. Arthritis Res Ther. 2007;9 Suppl 1:S1.

29. Hudson RS, Yi M, Esposito D, Watkins SK, Hurwitz AA, Yfantis HG, et al. MicroRNA-1 is a candidate tumor suppressor and prognostic marker in human prostate cancer. Nucleic Acids Res. 2012;40(8):3689–703.

30. Gilkes DM. Implications of Hypoxia in Breast Cancer Metastasis to Bone. Int J Mol Sci. 2016;17(10).

31. Keller ET, Brown J. Prostate cancer bone metastases promote both osteolytic and osteoblastic activity. J Cell Biochem. 2004;91(4):718–29.

32. Frieling JS, Shay G, Izumi V, Aherne ST, Saul RG, Budzevich M, et al. Matrix metalloproteinase processing of PTHrP yields a selective regulator of osteogenesis, PTHrP1-17. Oncogene. 2017;36(31):4498–507.

33. Ibrahim T, Flamini E, Mercatali L, Sacanna E, Serra P, Amadori D. Pathogenesis of osteoblastic bone metastases from prostate cancer. Cancer. 2010;116(6):1406–18.

34. Garcia AJ, Tom C, Guemes M, Polanco G, Mayorga ME, Wend K, et al. ERalpha signaling regulates MMP3 expression to induce FasL cleavage and osteoclast apoptosis. J Bone Miner Res. 2013;28(2):283–90.

35. Christine Mehner EM, Aziza Nassar, William R. Bamlet, Evette S. Radisky and Derek C. Radisky. Tumor cell expression of MMP3 as a prognostic factor for poor survival in pancreatic, pulmonary, and mammary carcinoma. Genes & Cancer. 2015;6:480–9.

36. Jie-Feng Huang W-XD, Jun-Jie Chen. Elevated expression of matrix metalloproteinase-3 in human osteosarcoma and its association with tumor metastasis. JBOUN. 2016;21(1):235–43.

37. Furic L, Rong L, Larsson O, Koumakpayi IH, Yoshida K, Brueschke A, et al. eIF4E phosphorylation promotes tumorigenesis and is associated with prostate cancer progression. Proc Natl Acad Sci U S A. 2010;107(32):14134–9.

38. Lynch CC, Hikosaka A, Acuff HB, Martin MD, Kawai N, Singh RK, et al. MMP-7 promotes prostate cancer-induced osteolysis via the solubilization of RANKL. Cancer Cell. 2005;7(5):485–96.

39. Sasaki K, Takagi M, Konttinen YT, Sasaki A, Tamaki Y, Ogino T, et al. Upregulation of matrix metalloproteinase (MMP)-1 and its activator MMP-3 of human osteoblast by uniaxial cyclic stimulation. J Biomed Mater Res B, Appl Biomater. 2007;80(2):491–8.

40. Kusano K, Miyaura C, Inada M, Tamura T, Ito A, Nagase H, et al. Regulation of matrix metalloproteinases (MMP-2,−3, −9, and −13) by interleukin-1 and interleukin-6 in mouse calvaria: association of MMP induction with bone resorption. Endocrinology. 1998;139(3):1338–45.

41. Ally MM, Hodkinson B, Meyer PW, Musenge E, Tikly M, Anderson R. Serum matrix metalloproteinase-3 in comparison with acute phase proteins as a marker of disease activity and radiographic damage in early rheumatoid arthritis. Mediators Inflamm. 2013;2013:183653.

42. Imamura Y, Sadar MD. Androgen receptor targeted therapies in castration-resistant prostate cancer: Bench to clinic. Int J Urol. 2016;23(8):654–65.

43. Su Q, Xin L. Notch signaling in prostate cancer: refining a therapeutic opportunity. Histol Histopathol. 2016;31(2):149–57.

44. Thalmann GN, Sikes RA, Wu TT, Degeorges A, Chang SM, Ozen M, et al. LNCaP progression model of human prostate cancer: androgen-independence and osseous metastasis. Prostate. 2000;44(2):91–103 Jul 1;44(2).

45. Frank SB, Schulz VV, Miranti CK. A streamlined method for the design and cloning of shRNAs into an optimized Dox-inducible lentiviral vector. BMC biotechnology. 2017;17(1):24.

46. Yu X AJ, Chun JH, Friedman AD, Heimfeld S, Cheng L, Civin CI. HES1 inhibits cycling of hematopoietic progenitor cells via DNA binding. Stem Cells. 2006;24(4):876–88.

47. Dang L YK, Wang M, Gaiano N. Notch3 signaling promotes radial glial/progenitor character in the mammalian telencephalon. Dev Neurosci. 2006;28(1-2):58–69.

48. Campeau E RV, Rodier F, Smith CL, Rahmberg BL, Fuss JO, Campisi J, Yaswen P, Cooper PK, Kaufman PD. A versatile viral system for expression and depletion of proteins in mammalian cells. PLoS One. 2009;4(8):e6529.

49. Li X, Sterling JA, Fan KH, Vessella RL, Shyr Y, Hayward SW, et al. Loss of TGF-beta responsiveness in prostate stromal cells alters chemokine levels and facilitates the development of mixed osteoblastic/osteolytic bone lesions. Mol Cancer Res. 2012;10(4):494–503.

50. Edick MJ, Tesfay L, Lamb LE, Knudsen BS, Miranti CK. Inhibition of integrin-mediated crosstalk with epidermal growth factor receptor/Erk or Src signaling pathways in autophagic prostate epithelial cells induces caspase-independent death. Molecular biology of the cell. 2007;18(7):2481–90.

