## supplementary data for "Notch3 Promotes Prostate Cancer-Induced Bone Lesion Development via MMP-3"

**FIGURE S1**

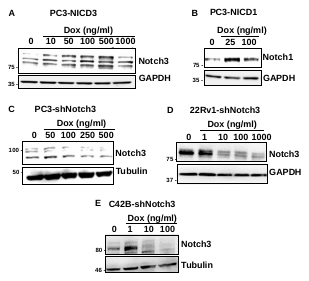

**Supplementary Figure S1. *In Vitro* induction of NICD3, NICD1, and knockdown of Notch3 in prostate cancer cells. (A)** Sub-confluent PC3 cells harboring Tet-inducible NICD3 (PC3-NICD3) or **(B)** Tet-inducible NICD1 (PC3-NICD1) were treated with increasing concentrations of Doxycycline (Dox) for 24hrs and cell lysates immunoblotted for Notch3 or Notch1. GAPDH served as loading control. Sub-confluent PC3 **(C)**, 22Rv1 **(D)** or C4-2B **(E)** cells harboring Notch3 shRNA (shNotch3) were treated with increasing concentrations of Doxycycline for 7 days and cell lysates immunoblotted for Notch3. Tubulin or GAPDH served as loading control.

**FIGURE S2**

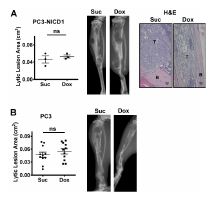

**Supplementary Figure S2. NICD1 or Doxycycline alone does not affect osteolytic lesion development.** **(A)** PC3 cells harboring Tet-induced NICD1 (PC3-NICD1) were injected into tibiae of 5-6 week old male NSG mice, treated with sucrose (Suc) and doxycycline (Dox) for 4 weeks, X-rayed, and lytic lesion area quantified. Representative X-ray images shown. Harvested tibiae were formalin fixed, decalcified, and stained with H&E. **(B)** Parental PC3 cells were injected into the tibiae of mice and treated with sucrose (Suc) or doxycycline (Dox) for 3 weeks, X-rayed, and lytic lesions quantified. Representative X-ray images shown. T = tumor; B = bone. Error bars are S.E.M, **(C)** n = 3, **(D)** n ≥ 11; ns = not significant.

**FIGURE S3**

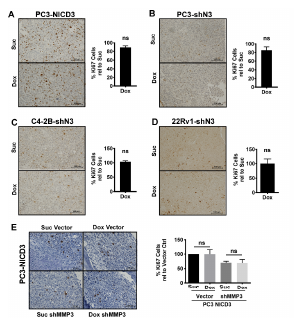

**Supplementary Figure S3. Notch3 does not alter tumor proliferation in the tibia.** Tibial tumors from mice injected with Tet-inducible **(A)** PC3-NICD3, **(B)** PC3-shN3 (Notch3 shRNA), **(C)** 22Rv1-shN3, **(D)** C4-2B-shN3, or **(E)** PC3-NICD3-vector or PC3-NICD3-shMMP3 were harvested from sucrose-treated (Suc) or doxyclycine-treated (Dox) mice and stained for Ki67 by IHC. Ki67 staining was quantified by counting 3-4 random fields at 20x magnification. Representative images shown. Error bars are S.E.M, n ≥ 7; ns = not significant.

**FIGURE S4**

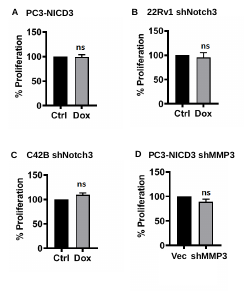

**Supplementary Figure S4. Notch3 not does alter proliferation of cells grown *in vitro*.** MTT assay was used to measure proliferation in **(A)** PC3-NICD3 cells treated with or without Dox (500ng/ml) for 1 day. **(B)**, 22Rv1 **(C)** or C4-2B cells harboring Notch3 shRNA (shNotch3) treated with Doxycycline (10ng/ml) for 7 days. **(D)** PC3-NICD3 cells harboring shMMP3 and its vector control. Error bars are S.E.M, n≥3; ns = not significant.

**FIGURE S5**

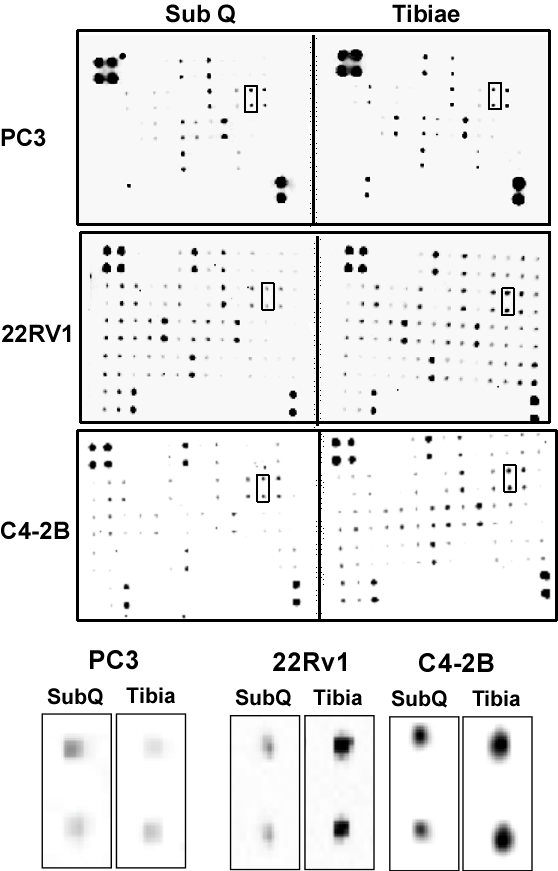

**Supplementary Figure S5. MMP-3 is upregulated in mixed/osteoblastic lesion tibiae compared to lytic tibiae.** PC3, 22Rv1, or C4-2B cells were injected subcutaneously (SubQ) or intra-tibially (Tibiae) and tumor lysates applied to a cytokine array. Levels of MMP-3 signal from the whole array (boxed pairs) were magnified and shown at the bottom.

**FIGURE S6**

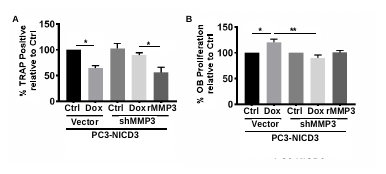

**Supplementary Figure S6. Recombinant MMP-3 rescues Osteoclast Phenotypes induced by shMMP3 cells. (A)** Bone marrow-derived osteoclasts or **(B)** osteoblasts differentiated in the presence of conditioned medium (CM) from doxycycline-treated (Dox) or vehicle-treated (Ctrl) PC3-NICD3-vector or PC3-NICD3-shMMP3 cells. PC3-NICD3-shMMP3 cells treated with recombinant MMP-3 (rMMP3) to rescue shMMP3 knock-down. **(A)** Percentage of TRAP+ cells quantified. **(B)** MTT assay of treated osteoblasts. Error bars are S.E.M, n≥3; *0.01≤ p≤0.05; **0.001≤p<0.01.

**TABLE S1: Antibodies**

**Target	Source	Product No	Blot	IHC**

**____________________________________________________________**

Notch1	CST	3608	1:1000

Notch3	CST	5276	1:1000

Notch3	Santa Cruz	5593	1:300

Notch3	Protein Tech	55114-1-AP	1:1000

MMP-3	Abcam	52915	1:75

Ki67	Thermo	RM-9106	1:100

Tubulin	Sigma	T9026	1:10,000

GAPDH	1:10,000

**TABLE S2: qRT-PCR Primers**

| Notch3-Human | Fwd | 5’CGTGGCTTCTTTCTACTGTGC |
| --- | --- | --- |
|  | Rev | 5’CGTTCACCGGATTTGTGTCAC |
| Notch1-Human | Fwd | 5’CGCAGATGCCAACATCCAGG |
|  | Rev | 5’ CCCAGGTCATCTACGGCGTTG |
| GAPDH-Human | Fwd | 5’GATCATCAGCAATGCCTCCTGC |
|  | Rev | 5’CTTCTGGGTGGCAGTGATGGC |
| MMP3-Human | Fwd | 5’ AGCAAGGACCTCGTTTTCATT |
|  | Rev | 5’ GTCAATCCCTGGAAAGTCTTCA |
| Bone Sialoprotein-Mus | Fwd | 5’ AAGCAGCACCGTTGAGTATGG |
|  | Rev | 5’ CCTTGTAGTAGCTGTATTCGTCCTC |
| ALP-Mus | Fwd | 5’ ATCTTTGGTCTGGCTCCCATG |
|  | Rev | 5’ TTTCCCGTTCACCGTCCAC |
| Osteocalcin-Mus | Fwd | 5’GCAATAAGGTAGTGAACAGACTCC |
|  | Rev | 5’ GTTTGTAGGCGGTCTTCAAGC |
| Osterix-Mus | Fwd | 5’ AGCGACCACTTGAGCAAACAT |
|  | Rev | 5’ GCGGCTGATTGGCTTCTTCT |
| Cathepsin K-Mus | Fwd | 5’TGACCACTGCCTTCCAATAC |
|  | Rev | 5’CTCTGTACCCTCTGCATTTAGC |
| Calcitonin Receptor-Mus | Fwd | 5’GACAACTGCTGGCTGAGTG |
|  | Rev | 5’ GAAGCAGTAGATAGTCGCCA |
| MMP9-Mus | Fwd | 5’ GATCCCCAGAGCGTCATTC |
|  | Rev | 5’ CCACCTTGTTCACCTCATTTTG |
| DC-STAMP-Mus | Fwd | 5’ GCGGAACTTAGACACAGGG |
|  | Rev | 5’ CAAAGCAACAGACTCCCAAATG |
| GAPDH-Mus | Fwd | 5’ GGAGAAACCTGCCAAGTATGA |
|  | Rev | 5’ TCCTCAGTGTAGCCCAAGA |
